# The mitoepigenome responds to stress, suggesting novel mito-nuclear interactions in vertebrates

**DOI:** 10.1101/2023.03.08.531655

**Authors:** John Lees, Fábio Pèrtille, Pia Løtvedt, Per Jensen, Carlos Guerrero Bosagna

## Abstract

The mitochondria are central in the cellular response to changing environmental conditions resulting from disease states, environmental exposures or normal physiological processes. Although the influences of environmental stressors upon the nuclear epigenome are well characterized, the existence and role of the mitochondrial epigenome remains contentious. Here, by quantifying the mitochondrial epigenomic response of pineal gland cells to circadian stress, we confirm the presence of extensive cytosine methylation within the mitochondrial genome. Furthermore, we identify distinct epigenetically plastic regions (mtDMRs) which vary in cytosinic methylation, primarily in a non CpG context, in response to stress and in a sex-specific manner. Motifs enriched in mtDMRs contain recognition sites for nuclear-derived DNA-binding factors (ATF4, HNF4A) important in the cellular metabolic stress response, which we found to be conserved across diverse vertebrate taxa. Together, these findings suggest a new layer of mito-nuclear interaction in which the nuclear metabolic stress response directly alters mitochondrial transcriptional dynamics.

## Introduction

Mitochondria are key mediators of the physiological and pathophysiological cellular responses to environmental stressors. The association between mitochondrial dysfunction and neurological disorders as well as cancer, Down syndrome, diabetes and infertility is becoming increasingly recognized (*1*, *2*). Importantly, mitochondrial dysfunctions are seldom linked to mitochondrial DNA (mtDNA) sequence alterations (*1*, *3*), fuelling a burgeoning interest in the mitoepigenome as the nexus linking environmental exposures, mitochondrial (dys)function and altered transcriptional dynamics.

Given the importance of retrograde and anterograde communication between mitochondrial and nuclear genomes, mitoepigenomic modification remains an unexplored mechanism by which stress can be embedded at the transcriptional level via the mitochondrial response (*4*). Certain environmental stressors and disease states are associated with alterations to the mitoepigenome (*5*–*7*) and have been used to investigate the extent and role of mitoepigenomic modifications, primarily in an in vitro context. Targeted bisulphite sequencing indicates that the mitochondrial D-loop control region is often differentially methylated upon exposure to, for example, airborne particulate matter (*8*). An in vitro study using MeDIP-Seq to interrogate the entire mitochondrial genome, revealed that 7 mitochondrial genes are hypomethylated in hepatocytes exposed to valproic acid, including MT-CO1 and MT-CO2 (*9*). At a whole organism level, there are a limited number of in vivo studies investigating mitoepigenomic response to environmental exposures. These studies often investigate a subset of mitochondrial genes, precluding a complete quantification of the mitoepigenetic response (*8*, *10*, *11*). In addition to the paucity of data regarding chemical exposures and disease, it remains to be known whether non-chemical environmental stressors are able to affect the mitoepigenome. For example, it is currently unclear whether circadian disruption, a known affecter of metabolism, influences the mitochondrial epigenome.

Circadian stress is an important model in which to investigate the mitoepigenomic response to stress. Circadian rhythmicity is essential in driving the appropriate temporal patterns of metabolism, meaning disruption of circadian rhythmicity results in aberrant metabolism (*12*). Successful circadian rhythmicity requires a ‘zeitgeber’ to synchronise endogenous clocks, primarily in the form of photic input. In mammals the pineal gland is under the control of the suprachiasmatic nucleus (SCN) in the hypothalamus, which is affected by light exposures (*13*). In birds, however, photic input occurs both through the retina and the pineal gland which functions as an independent oscillator and, together with the SCN, directs rhythmicity in behaviour, physiology and immunity in response to ambient light conditions (*14*, *15*). The pineal gland’s function lies primarily in the circadian secretion of melatonin, whose circulating levels vary in response to photoperiod. Deviation of light conditions from those required to generate a normal daily rhythm may disrupt the core circadian machinery, including the melatonin rhythmicity, leading to suboptimal physiological states (*16*–*18*). Interestingly, melatonin is intimately linked to mitochondrial function, whose synthesis within the mitochondria may have arisen early in the endosymbiotic origins of eukaryotes (*19*). As such, melatonin is thought to play a neuroprotective role, with circulating levels representative of age and susceptibility to psychiatric disorders. Indeed, melatonin’s protective action acting through mitochondria may provide an ameliorating role in the development of cancer cells (*20*). Importantly, circadian stress is associated with alterations to methylation of the genomic DNA (*21*). Whether such methylation changes are mirrored in the cell’s mitochondrial genome is yet to be determined.

Here we show that random illumination patterns elicit mito-epigenomic effects within the pineal gland and identify epigenetically plastic regions (mtDMRs) which respond to stress in a sex-specific manner. We make use of a robust MeDIP-Seq approach that controls for the false detection of mitochondrial methylation resulting from nuclear mitochondrial pseudogenes. Using this method, we reveal the extent of mitochondrial cytosine methylation throughout the mitochondrial genome in the pineal glands of both male and female chickens (*Gallus gallus domesticus,* Linneaus) in response to externally induced circadian stress, taking advantage of the fact that avian pineal glands respond directly to photic stimuli.

## Results

### Sequencing and alignment

Sequencing of the 12 MeDIP-seq libraries relating to control and circadian stressed male and females gave an average of approximately 2.9 × 10^5^ of reads assigned to the reference chromosome MT (*Gallus gallus 6.0,* NCBI). Upon controlling for nuclear mitochondrial pseudogenes, we mapped 2.7 × 10^5^ reads, which corresponds to 2.2±0.5 × 10^4^ reads per individual, this corresponds to 93.6% of the reads mapped to the chrMT prior to correction. The average depth of these aligned reads was 172.6 X ± 39.2 X and covering 99.9 % (±0.1) of the chicken mtDNA genome. On average, the alignment against the mtDNA presented 32.6 times more depth of coverage than the whole chicken reference genome (Table S1).

### Identification of mtDMRs

Four pairwise comparisons were performed on the mtDNA sequencing data to identify differentially methylated CpN dinucleotides (DMCpNs) resulting from circadian stress and as a result of sex; Male control vs. female control (MC vs. FC), male stress vs. female stress (MS vs. FS), male control vs. male stress (MC vs. MS) and female control vs. female stress (FC vs. FS). We identified 62 significant mtDMRs across all statistical comparisons (Table S2), 46 of which had a length greater than 5bp (range= 5-387bp). 15 of these mtDMRs were present in at least two of the pairwise statistical comparisons, while 47 were unique regions within each contrast (Figure 1). Base composition within the mtDMRs did not differ significantly from the expected proportions within the mitochondrial genome, with the exception of G, which differed from the expected frequency in our MS vs. FS comparison (p=0.01, Table S3). Methyl cytosines within the mtDMRs showed differential coverage in all four possible CpN dinucleotide combinations (table S4). The number of CpNs exhibiting differential coverage was not significantly different from the total number of each CpNs within a particular mtDMR region (Table S4). Furthermore, the proportions of CpNs within mtDMRs were not different from the expected proportions within the whole mitochondrial genome (Table S4). Although suggestive of differential cytosine methylation, differential coverage in MeDIP-Seq does not allow for the definitive identification of cytosine methylation with single base accuracy. It is therefore not possible to identify whether certain CpNs are more commonly differentially methylated than others based on differential coverage alone. With this in mind, an alternative way to determine the importance of different CpNs within the mtDNA is to look at the frequency of mtDMRs containing no instances of a particular CpN. In this case the absent CpN cannot be driving the significant differences in coverage, meaning that the differential coverage in these mtDMRs is a result of differential methylation of the other three CpNs. In this context, we found that CpG motifs were absent in 43.5% of the mtDMRs, whereas the other CpNs showed lower absences (CpA = 22.6, CpT = 17.7, CpC = 21).

**Fig 1.**
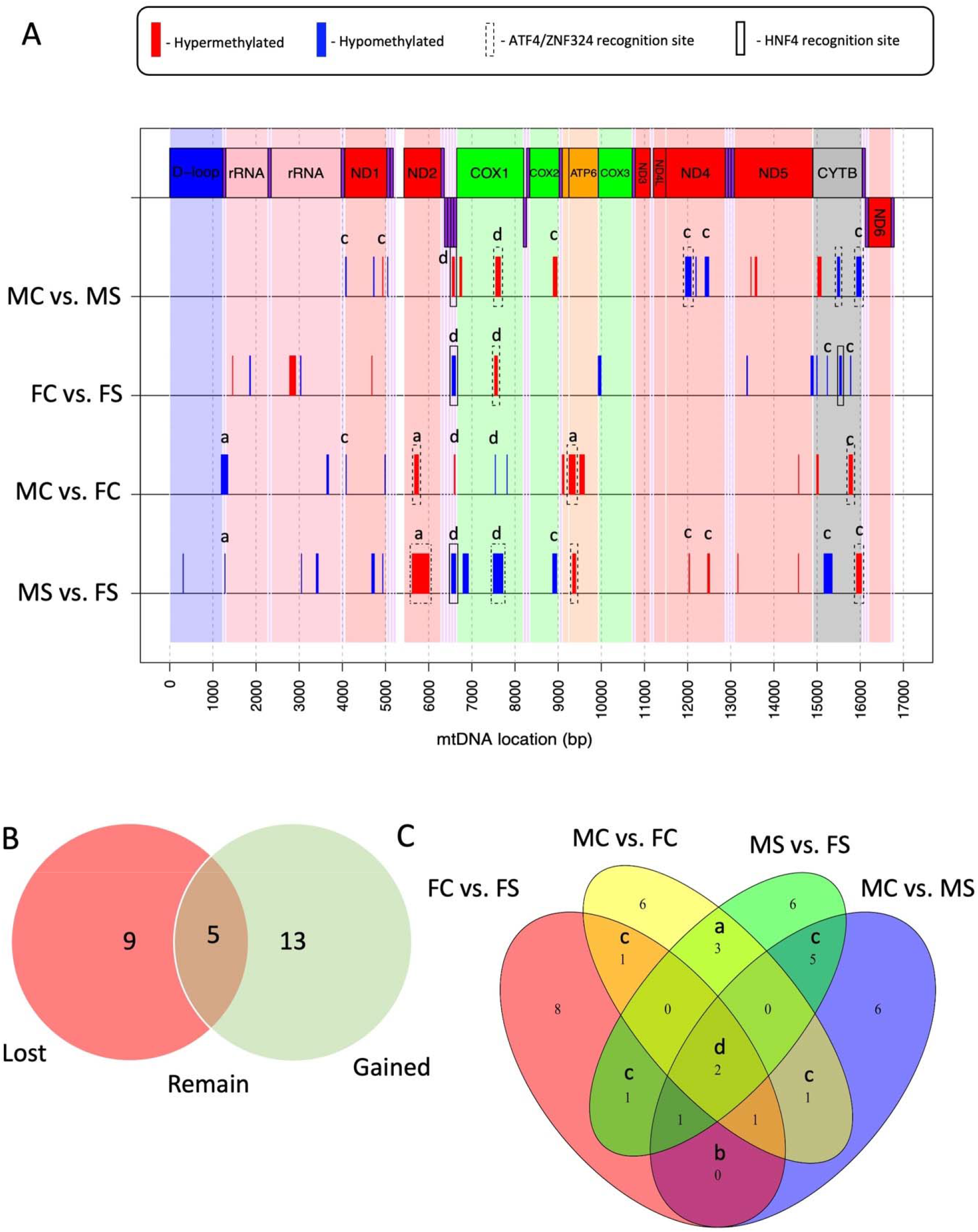
The numbers and locations of regions of differential methylation (mtDMRs) in response to sex and stress. **(A)** The location of regions of differential cytosine methylation (mtDMRs) across the mitochondrial genome of pineal gland mitochondria resulting from circadian disruption (Stress). Within the 4 pairwise statistical comparisons, regions of hypo (blue) and hyper (red) methylation relate to our pairwise statistical comparisons; Male control vs. female control (MC vs. FC), male stress vs. female stress (MS vs. FS), male control vs. male stress (MC vs. MS) and female control vs. female stress (FC vs. FS). Coding regions within the heavy strand (mtDNA genes, above line) and light strand (mtDNA genes, above line) of the mitochondrial DNA are displayed for reference, with subunits of the same electron transport complexes shown in the same colour. tRNAs are shown in purple. Boxes around regions of differential methylation indicate the presence of sequence motifs associated with HNF4 (solid box), ATF4/ZNF324 (dashed box) or both (dot and dashed box). Letters within each window represent the nature of the mtDMR. a = sex-specific mtDMRs unrelated to stress, b = mtDMRs with identical responses in males and females, c = mtDMRs differing in the control and stress of one sex only, d =mtDMRs different across all statistical comparisons. **(B)** Venn diagram showing the mtDMRs which are lost, gained or remain between MC vs. FC and MS vs. FS, representing sex-specific stress-related regions (gained/lost) and sex-specific, stress-independent regions (remained). **(C)** A Venn diagram showing the numbers of mtDMRs which are either unique or present across our pairwise statistical comparisons.

### Sex differences in mtDMRs

A comparison of mtDMRs present in the MC vs. FC with those present in MS vs. FS revealed a predominant increase in the number of mtDMRs upon exposure to circadian stress. A total of 9 mtDMRs, which were present in MC vs. FC comparison, were lost upon exposure to circadian stress (i.e. no longer present in the MS vs. FS) whilst 13 were gained, representing sex-specific stress-related regions (gained/lost). In comparison, five mtDMR sites remained, representing sex-specific, stress-independent regions (Figure 1). Of the total mtDMRs discovered, 25% were represented in at least two of the four contrasts. The extent of reoccurrence of mtDMRs in different contrasts allowed for the identification of the nature of their response to both stress and sex (Figure 1). We found that the most commonly recurring mtDMRs were those identified in both MS vs. FS and MS vs. MC comparisons (i.e. ‘Male stress-specific regions’, of which there were five mtDMRs), those differentially methylated only as a result of sex (i.e. ‘sex-specific regions’, which we found in both MC vs. FC and MS vs. FS comparisons, of which there were three mtDMRs) and regions which were differentially methylated as the result of sex and stressor (‘broadly-affected regions’, which were present in all statistical comparisons, of which there were two mtDMRs).

### Hypo vs. hypermethylation

Within the mtDMRs, the direction of methylation was strongly influenced by stress and sex. Overall, mtDMRs were hypomethylated in FC in comparison to FS birds (10:4 hypo-:hyper-methylated mtDMRs in FC compared to FS) whereas MC vs. MS show equal levels of hyper and hypomethylation (Figure 1). The mtDMRs showed similar levels of hypo- and hyper-methylation when comparing MC vs. FC birds whereas MS birds showed a larger proportion of hypo-methylated regions compared to FS individuals (11:7 hypo-:hyper-methylated mtDMRs in MS compared to FS birds).

#### Motif discovery and enrichment analysis

Within mtDMRs, five significant motifs were found within the mtDMRs (XSTREME, E < 0.05, minimum width 6 bp, table 1, figure 1). Through a comparison of these motifs with three vertebrate motif databases, two of the motifs were found to share composition with known vertebrate motifs: The SGACTSTG motif was associated with the binding of hepatocyte nuclear factor alpha and gamma (HNF4A/HNF4G) and the CACCATCCT motif was associated with the binding of both zinc finger protein 324 (ZNF324) and the activating transcription factor 4 (ATF4). HNF4A/HNF4G binding-domains were identified in four mtDMRs located in all but the MC vs. FC comparison. Three of these were present in a single region from bases 6531-6625 (coding for tRNA-Cys and tRNA-Tyr) and were differentially methylated in MC vs. MS, FC vs. FS and MS vs. FS comparisons. Within this region, the pattern of methylation after stress differed in direction between males and females, being hypomethylated in stressed males (MS) in comparison to control males (MC) and hypermethylated in stressed females (FS) in comparison to their control counterparts (FC). Additionally, twelve ZNF324/ATF4 motifs were identified within mtDMRs and were found across all comparisons. Of these twelve sites, nine occurred within just 4 mtDMRs located within the *ND2, COX1, ATP6* and *CYTB* coding regions. The broadly affected *COX1* region (bp 7499-7717) showed the same pattern of hypomethylation in stressed males and females compared to their relative controls, although basal and stressed levels of methylation in females were higher than those of equivalent males. Other ATF4 sites commonly showed a pattern in which basal levels of methylation were unaffected by stress but were hypomethylated in females in comparison to males.

**Table 1.** Consensus motifs identified within mtDMRs of the mtDNA of chickens exposed to circadian stress. All significance values for the Homologous known vertebrate motifs identified through a comparison with vertebrate motif databases are shown where applicable. P values were generated using XSTREME (Motif Discovery and Enrichment Analysis): Version 5.4.1 released on Sat Aug 21 19:23:23 2021 −0700

| Rank | Consensus sequence | Width (bp) | Number of sites within mtDMRs | Sea P value | E value | Homologous motif |
| --- | --- | --- | --- | --- | --- | --- |
| 1 | SGACTSTG | 8 | 14 | <0.001 | <0.001 | HNF4G, HNF4A |
| 2 | AYWAKAA | 7 | 16 | <0.001 | <0.0001 | NA |
| 3 | ATTATGAA | 8 | 13 | <0.001 | <0.0001 | NA |
| 4 | ACWACAGCC | 9 | 11 | <0.001 | <0.001 | NA |
| 5 | CACCATCCT | 9 | 17 | <0.001 | <0.0001 | ATF4 DBD, ZNF324 |

#### Motif presence across diverse taxa

Given the highly conserved nature of the mitochondrial genome across eukaryotes, we might expect that functionally important motifs within the mtDNA are conserved across diverse taxa. Having identified DNA-binding motifs (ATF4, ZNF324 and HNF4A) enriched in mtDMRs, we therefore sought to establish the prominence of these motifs within mitochondrial genomes of chickens (*Gallus gallus*), humans (*Homo sapiens*), alligators (*Alligator mississippiensis*), zebrafish (*Danio rerio*) and fruit flies (*Drosophila melanogaster*). Binding motifs were taken from human databases (tables S5, S6 and S7) and compared against the mitochondrial genomes using the FIMO web-based tool (*22*).

### Identification of ATF4 recognition sites

Five instances of the ATF4 binding motif were identified within the *G. gallus* genome, which were located in three protein-coding regions (16S rRNA, COX3 and ND4), while there were between two and four ATF4 sites present in the other species tested (*H. sapiens* (3), *A. mississippiensis* (2), *D. rerio* (4), and *D. melanogaster* (3)), with 76% of sites occurring on the mitochondrial light (negative) strand (Figure 2, Table S8). Two protein-coding regions, 16S ribosomal rRNA and ND4 contained 53% of these sites and one region was found within the 16S ribosomal RNA gene in all species except for *D. melanogaster.* Alignment of the common ATF4 binding domains within the 16s RNA sequences (± 20 bp upstream and downstream) revealed 100% conservation between species in the core binding site sequence (TGTTGGATCAGGAC) and a 70% consensus sequence differing in only 2/54 bases.

**Fig 2.**
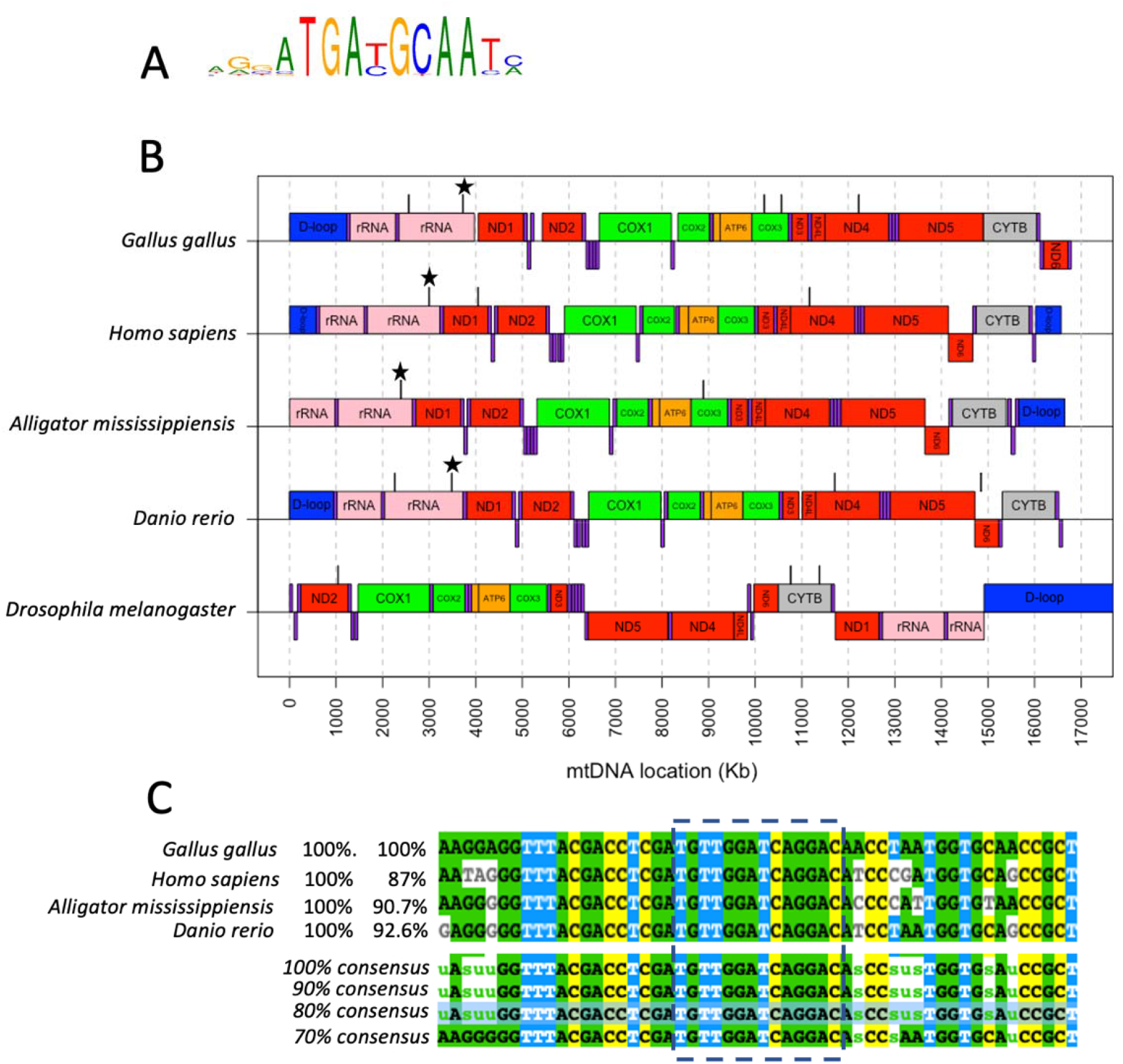
ATF4 presence in mtDNA across diverse taxa. **(A)** The DNA recognition motif for human ATF4 transcription factor used for a comparison across diverse taxa. **(B)** Locations of human ATF4 transcription factor recognition sites (black bars above mtDNA coding regions) identified using FIMO) within the mitochondrial genomes of diverse taxa: *Gallus gallus, Homo sapiens, Alligator mississippiensis, Danio rerio* and *Drosophila melanogaster*. Coding regions within the heavy strand (mtDNA genes, above line) and light strand (mtDNA genes, above line) of the mitochondrial DNA are displayed for reference, with subunits of the same electron transport complexes shown in the same colour. tRNAs are shown in purple. **(C)** Sequence allignment of the 16S rRNA region associated with ATF binding (shown with star in b, outlined in dashes in c) as well as 20bp upstream and downstream of the site.

### Identification of ZNF324 recognition sites

The chicken mitochondrial genome contained 10 sites sharing sequence with that of the ZNF324 recognition site (Figure S1, Table S9). Despite being found in all species tested, fewer sites were located in the mitochondrial genomes of *H. sapiens* (6), *A. mississippiensis* (8), *D. rerio* (5), and *D. melanogaster* (2). Recognition sites were found in the ND2, ND4 and ND5 genes in three of the five species, each coding for components of electron transport chain complex I. 94% of the recognition sites were located within the mitochondrial light (negative) strand.

### Identification of HNF4A recognition sites

Our analysis revealed five regions of the *G. gallus* mtDNA containing sequences matching the HNF4A recognition domain (Figure S1, Table S10), with fewer sites found in *H. sapiens* (4), *A. mississippiensis* (2), *D. rerio* (1), and *D. melanogaster* (0). Unlike ZNF324 and ATF4 sites, location of the HNF4 recognition sites was variable across species with no apparent commonalities in location. 92% of sites were located on the mitochondrial heavy (positive) strand.

## Discussion

With few exceptions, both wild and domesticated species have evolved under the influence of circadian fluctuations in light intensity. Disruption of the resultant evolved circadian rhythmicity represents a form of environmental stress that is relevant in multiple contexts from animal welfare to human health and ecology (*23*, *24*). Our results demonstrate that the mitoepigenome of the pineal gland, a core regulator of circadian rhythmicity in vertebrates, is impacted by circadian stress during postnatal development when deviations from a 12:12 light dark cycle occur early in life. Importantly, the nature of this epigenetic response differs between the sexes.

### The effects of circadian stress upon mtDNA methylation

The notion of mtDNA methylation remains controversial, with numerous studies reporting an absence or minimal level of methylation (*25*–*28*). Our data clearly demonstrate that in the pineal gland of chickens not only is mtDNA methylated, but cytosinic methylation of mtDNA is variable upon developmental exposure of animals to circadian stress. Moreover, the effects are different between the sexes. The novel finding that mtDNA within the pineal gland is extensively methylated, often in non-CpG contexts within distinct epigenetically plastic regions (mtDMRs) of the mitochondrial genome has stark implications for our understanding of mitochondrial function and malfunction. Although circadian disruption is known to have metabolic effects, this study provides the first evidence that an environmental stressor, as opposed to a disease state (*2*, *6*) or chemical exposure (*8*–*11*), is capable of modifying the epigenetic landscape of the mitochondria. Additionally, the identification of transcription factor binding motifs within the mtDMRs identified adds evidence to the notion (*29*) that nuclear-derived transcription factors play a role in regulating mtDNA transcription across species.

### Locations of mtDMRs

Although mtDMRs were found throughout the mitochondrial genome, across all statistical comparison, mtDMRs were biased towards specific coding regions of the mtDNA. For example, *CYTB, ND1, COX1 and ND5* contained over 50% of all the detected mtDMRs (figure 1). All four regions are protein coding genes for key components of electron transport chain respiratory complexes I (*ND2, ND4*), complex III (*CYTB*) and IV (*COX1*). Altered transcriptional activity of these genes would have clear functional consequences for electron transport, proton gradient formation and thus the ATP production efficiency of the affected mitochondria. Overall, the finding that specific regions of the mtDNA are more responsive than others lends strong evidence to the notion that the mitoepigenome is regulated and responsive to the environment in a targeted manner. The observation that the CpN composition within mtDMRs is not different from that of the mitochondrial genome as a whole adds weight to this notion, suggesting specific motifs may be important in driving mtDMR methylation as opposed to cytosine content, as is the case in CpG islands within the nucleus (*30*).

Certain genes within the mtDNA contained very few to no mtDMRs, indicating a lack of cytosinic methylation. Of particular interest is the finding that cytosine methylation of the D-loop control region is not responsive to stress. The D-loop is central to mitochondrial transcription and mtDNA replication, representing the origin of transcription around the DNA plasmid. Because of its importance in transcriptional dynamics, previous studies have often focused on the D-loop in order to detect methylation. Alterations to D-loop methylation have been previously observed in response to environmental stress (*10*) (*8*) (*11*, *31*) and disease (*32*, *33*) (*34*) (*35*). The absence of D-loop mtDMRs here may indicate that this region is less responsive to circadian stress. Alternatively, it is possible that individual cytosines at specific locations within the D-loop are important but were missed in our analysis, which was biased towards regions of differential methylation spanning multiple cytosine bases.

### Sex differences in mtDMRs

Our data reveal a clear sex difference in the mitoepigenome, both in terms of baseline methylation levels between control male and female birds and in the response of the pineal gland to circadian stress. By comparing the differential methylation of mtDMRs between control and stressed male and females, we were able to identify regions of the mtDNA associated with sex, stress and the interaction of the two. Control males and females differed in mtDNA methylation within one tRNA, 2 rRNA and 7 protein-coding genes, 3 of which coded for sub-units of electron transport chain complex 1 (ND1, ND2 and ND5). Interestingly mtDMRs within *atp6, trna1* and *nd2* remained differentially methylated between both stressed and unstressed males and females, resenting sex-related epigenetic elements. Furthermore, two of these ‘sex-specific regions’ contained binding sites for the ATF4 transcription factor, hinting at a basal sex difference in mito-nuclear interactions. These findings are concomitant with findings showing sex differences in metabolism and mitochondrial function throughout the animal kingdom, with both animal and human studies demonstrating divergence in the stress responses of males and females (*36*–*38*).

Previous studies in pigs indicate opposite directionality of mtDNA promoter methylation and transcription between males and females in response to stress (*31*). Looking at the entire mitochondrial genome, we find elements in which mtDNA which respond both oppositely (e.g. *trna10,* which is hypomethylated in stressed males and hypermethylated in stressed females) and similarly (*cox1,* which is hypomethylated in stressed males and females), yielding a more complex picture of the mitochondrial methylome. Interestingly, the *trna10* site that responds oppositely contains a motif recognition site for Hnf4 stress-related transcription factor whereas the *cox1* mtDMR contains an Atf4 stress-related transcription factor binding site. Should these transcription factors bind differentially based on methylation state, these results hint at a complex picture in which different nuclear-derived transcription factors exert differential effects based both upon sex and the location within the mtDNA.

### Cytosine context within mtDMRs

In relation to the nucleotides adjacent to methyl-cytosines within the mtDNA, our results agree with previous findings from diverse taxa such as mice (*39*) and fish (*40*), suggesting that non-CpG methylation is relevant and widespread in vertebrates. Investigations utilizing NGS with primarily mammalian models have detected higher non-CpG vs CpG methylation within the mtDNA (e.g. (*39*, *41*)). Non-CpG methylation is more commonly observed in prokaryotes, however is also relevant to eukaryotic genomic DNA (despite the predominance of CpG methylation) and is increasingly being considered as relevant to transcriptional processes within the nucleus (*41*, *42*). As opposed to CpG islands found within the nuclear genome, significant regions of differential methylation within the mtDNA contained methylated cytosines within CpN motifs in similar proportions to those within the mtDNA as a whole. Hence, the location of the methylated region within specific mtDNA motifs may be more critical to any functionality or DNA response than simply CpN bases being methylated per se. Methylated cytosines are more likely to transition to a T during DNA replication of the DNA. As a result of the predominance of methyl cytosines in a CpG context within the nucleus, it is perhaps not surprising that across the animal kingdom, age-related C>T transitions within the genomic DNA occur primarily in CG dinucleotides (*43*). Interestingly, the data of Cagan et al., 2022 reveal that C>T transitions in mtDNA with age occur commonly within a CpT context (particularly at GCT motifs), in keeping with our data showing extensive CpT representation within regions of differential methylation. mtDNA methylation may therefore not be only of interest in a mechanistic context but also in an evolutionary context, in which C>T transitions within certain regions may have shaped the mitochondrial genome through time (*44*). It would therefore be of interest to determine whether C>T transitions within the mtDNA are located within distinct regions across genera.

### Mechanistic consequences of differential methylation

Having discovered extensive cytosine methylation within the mtDNA, it is important to establish the potential mechanistic consequences of differential methylation within the mtDNA. By identifying functional regions which are plastic in their epigenetic response, we suggest that differential cytosine methylation within the mtDNA is indicative of functionality. Here, mtDMRs responding to circadian stress contained binding motifs associated with nuclear-derived transcription factors known to be central to metabolic homeostasis and the cellular stress responses (HNF4, ATF4 and ZNF324). The importance of these transcription factors in modulating stress-responses coupled with the increasing evidence that nuclear transcription factors locate to the mitochondria (*29*) add weight to the novel idea that nuclear stress signals directly alter mitochondrial gene transcription through binding of DNA elements throughout the mitochondrial genome. It appears that mtDNA transcription is therefore driven by more than simply the mitochondria-specific POLRMT polymerase and the TFAM and TFB2M transcription factors thought to dominate mtDNA transcription.

Among the transcription factors found to have binding motifs within mtDMRs, ATF4 is a transcription factor at the core of the mitochondrial unfolded protein stress response and integrated stress response (*45*). Furthermore, ATF4 is correlated with mitochondrial function in non-stressed cells, suggesting a role in not only the mitochondrial stress response, but in normal mitochondrial homeostasis (*46*). Exposure of HeLa cells to 4 differing mitochondrial stressors revealed that ATF4 binding sites were present in many of the genes involved in the varying stress responses (*46*). Crucially, mitochondrial stress was found to inhibit mitochondrial translation and decrease mitochondrial ribosomal proteins in what the authors state as a ‘localized response’. Our findings of a highly conserved region across species within the 16s ribosomal DNA of the mitochondria lends evidence to the idea that ATF4 may, in fact, be mechanistically linked to the alterations in translation observed in stressed mitochondria as opposed to part of a separate response.

Transcription factor binding specificities are largely conserved across the bilateria (*42*, *47*, *48*)and the roles of ATF4, HNF4 and zinc finger proteins as regulators of metabolism, as well as their expression patterns, are highly conserved from flies to mammals (*49*). Therefore given our observation of recognition sites for these DNA-binding proteins across a range of metazoan species, it is reasonable to infer that mitochondrial roles of these nuclear transcription factors may also have evolved early in animal evolution. In addition, the identification of binding sites through mtDMRs suggest that methylation may be an important regulator of this transcription factor binding, a known phenomenon regarding protein-DNA interactions (*50*, *51*) for example involving zinc-finger proteins (*52*). Importantly, the presence of extensive non-CpG methylation within genomic DNA (*53*, *54*) in addition to the observation that CpA methylation is important in altering DNA binding affinity (*51*, *52*) together suggest an important role of non-CpG methylation within the nucleus as well as within the mitochondria.

As well as identifying motifs consistent with DNA-binding factors, our mtDMRs reveal additional novel motifs with no clear eukaryotic homologues. Given that, unlike the nuclear DNA, mtDNA is bacterial in origin, it should also be considered that mtDNA transcription could additionally be regulated in a ‘non-nuclear’ way. Bacterial plasmids are actively methylated and regulatory elements such as enhancers are found in bacteria (Xu and hoover, 2001). The mtDMRs detected without eukaryotic equivalents may therefore represent important motifs involved in an, as yet, undetermined process of mtDNA regulation.

Understandably, the close link between transcription and methylation within the nucleus bias explanations of mtDNA methylation towards transcription factor binding. Despite this, there are alternative functional explanations. DNA methylation, through the association with methylated DNA-binding proteins, can influence the rate of polymerase translocation across the DNA. mtDMRs could thus represent pause or slow sites across the mtDNA akin to those important for gene regulation in bacteria (*55*). Methylation of the mtDNA could also be involved in altering DNA stability and 3-dimensional structure. In the absence of histone proteins, mtDNA accessibility can be altered by the 3-dimensional structure of the nucleoid, altering transcriptional dynamics. It is realistic to hypothesize that mtDNA methylation could influence the conformation of the mtDNA or, by altering the affinity of DNA binding proteins such as TFAM, could influence DNA packaging and flexibility (*56*, *57*).

## Conclusions

In conclusion, our results demonstrate that the mitoepigenome within pineal cells of chickens is dynamic and responsive to circadian stress, adding weight to the burgeoning evidence that mtDNA is methylated in a systematic and dynamic manner. Furthermore, we have identified regions of differential methylation within the mtDNA that appear to have a role in the nuclear-driven regulation of mitochondrial transcription in response to environmental stress. Although it is not possible to determine to what extent mtDNA methylation represents a causative or correlative element, these data add a novel layer of complexity to our understanding of the subcellular response to circadian disruption. The widespread presence of mtDMRs, some which are associated with known stress-related nuclear transcription factors suggest a nuanced mitochondrial genome response to circadian stress involving more than just the three transcription factors currently thought to control the mtDNA transcriptional responses. In order to unpick this response, it is essential that future experiments aim to connect mtDNA methylation, transcription factor binding and differential transcription of the mitochondrial and nuclear genomes.

## Materials and Methods

### Animals and husbandry

Fertilized Hy-Line (Swedfarm, Linghem, Sweden) white leghorn chicken (*Gallus gallus domesticus*) eggs were incubated in a 60% humidified atmosphere at 37.8°C with rotation once every hour, until hatching. After hatching, all individuals were wing tagged, placed in boxes, and directly transferred to cages. They lived in mixed groups containing both sexes (females n=18, males n=16), and between 5 and 6 chickens per cage. They were not confidently sexed until culling and were therefore not balanced between groups. Throughout the experiment all birds had *ad libitum* access to water and food as well as wood chips on the floor of every cage. Feed trays including starter food provided access to food within the first days, which was later on replaced by hanging food (Pennafood). During the first period of the experiment, chickens had access to heated roofs. The animals were raised in temperature and light-emitting diode (LED) light-controlled cages. The handling and maintenance of chickens used in this study were conducted under the permission of the Regional Committee for Scientific Research on Animals, (License Nr. 50-13 Linköping).

### Treatment

The animals were randomly divided into two groups after one day of incubation. The control group (n=6, 3 males and 3 females) was kept at standard 12:12 light-dark cycle for their entire life. The circadian stress group (n=6, 3 males and 3 females) was exposed to continuous light from days 3-6 of age. From days 7-24, birds experienced a randomized light schedule (36h of light every 3 days delivered at random intervals between 3 to 21 hours). Over a three-day period, the total amount of light and dark was equal to that of control animals. From day 25 and for the rest of the experiment, the animals were kept at standard 12:12 light-dark cycle.

Each cage received light from two LED lamps, controlled by a timer. Cages were isolated to stop light from reaching neighbouring cages, confirmed by light measurements before the experiments. Culling took place over a three-day period when the birds were 34-36 days old. Control and circadian-stressed chicks were sacrificed by decapitation and pineal glands were removed from the brain and snap frozen in liquid nitrogen.

### DNA isolation

In order to isolate total DNA from pineal gland tissue, Allprep®DNA/RNA/miRNA Universal kit (Qiagen GmbH, Germany) was used. First, the complete pineal gland was diluted in 350μl RLT Plus buffer, which contained guanidine thiocyanate for tissue lysis and then homogenized for 40 seconds in a rotor-stator homogenizer (FastPrep^-24^ Homogenizer, MP^™^ Biomedicals, California, USA). Further steps were performed according to the manufactures protocol. Samples was stored at −80°C. The DNA concentration was determined by nanodrop spectrophotometer and calculated by Gene Quant pro RNA/DNA calculator Software (NanoDrop ND^-1000^ Spectrophotometer, Thermo Scientific, Wilmington, USA).

### Ion torrent MeDIP-Seq library preparation

MeDIP-Seq libraries for Ion Torrent semiconductor sequencing were performed on the Ion Torrent PGM using a modified protocol from Ion Plus Fragment Library kit (Catalog # 4471252) combined with a 5-hydroxymethylcytosine (5hmC Kit, Catalog # AF-110–0016) immunoprecipitation kit (Diagenode) according to a previously optimized protocol (Guerrero-Bosagna & Jensen, 2015). For 5mC MeDIP-Seq, 8 μg of DNA was sheared to a mean fragment size of 350 bp by sonication in a Fisher ultra-sonicator, with probe attached to a cooling cup horn. The final preparation of the libraries was performed at the SNP&SEQ facilities of the SciLifeLab (Sweden) and consisted of the end-repair, purification with Agencourt AMPure XP reagent beads (Beckman Coulter), and ligation with Ion Torrent compatible sequencing adapters following the standard Ion Plus Fragment Library Kit protocol. Adapter-ligated DNA was purified using Agencourt AMPure XP reagent beads and immunoprecipitated using a 5-methylcytosine kit (MagMeDIP) according to the manufacturer’s protocol. Following 5mC MeDIP, libraries were size-selected from 200–400 bp on a 2% agarose E-gel (Invitrogen) and extracted using a MinElute Gel DNA extraction kit (Qiagen) according to the manufacturer’s instructions. Libraries were then amplified at 18 cycles of PCR (95°C, 15 s; 58°C, 15 s; 70°C, 1 min), purified using MinElute columns (Qiagen), and eluted in 16 μl EB (Qiagen). Libraries were assessed for quality and concentration on an Agilent Bioanalyzer instrument.

### Ion torrent sequencing

MeDIP libraries were templated using the Ion OneTouch 2 System instrument with either the Ion PGM Template OT2 200 kit for the PGM instrument or the Ion PI Template OT2 Kit v2 for the Proton instrument according to manufacturer’s protocols (Life Technologies). Following template reactions, templated libraries were assessed for quality using the Ion Sphere Quality Control Kit (Catalog # 4468656) and Qubit fluorometer (Life Technologies). Templated libraries that passed manufacturer recommended criteria were enriched using the Ion OneTouch ES instrument. Immediately after enrichment, MeDIP libraries were sequenced on either the PGM with 318 chips or Ion Proton with the P1 chips using the Ion PGM 200 Sequencing kit or Ion PtdIns Sequencing 200 Kit v2, respectively, according to instructions provided by the manufacturer (Life Technologies).

### MeDIP-Seq analysis

Ion Torrent sequencing reads were quality-filtered and aligned to the chicken reference sequence *Gallus*_*gallus* 6.0 (NCBI) using Torrent Suite Software’s (v4.0) TMAP alignment software. Alignment files were processed into the BAM format and used for further analyses. The index and coverage depth of the “.bam” files were performed using Samtools v.1.11 (*58*) with the “index and “depth” options.

To control for nuclear-mitochondrial pseudogenes within Methylated DNA Immunoprecipitation Sequencing data, we used a recently published pipeline (*59*), which consists in four steps using the previously mentioned BAM files as input. First, we extracted the new BAM file of “unique reads”, which were aligned to no more than one specific region of the reference genome (RG). For that, we used Samtools v.1.11. Second, we extracted a FASTQ file with those “unique reads” from the previous generated BAM files using BEDTools v.2.29.2 to be aligned against an RG index without the chrMT. This reference genome was build using “bowtie2-build” function from Bowtie v.2-2.4.2. Third, we extracted a new BAM file containing the non-aligned reads from the previous BAM file generated by the last step. Forth, we extract the .fasta file from the bam file of the earlier step to be aligned to a new genome index, but now build using only the mtDNA sequence from the RG.

Through these last BAM files generated from the previous pipeline, the MEDIPS R (Lienhard et al., 2014) package was used for basic processing, quality controls, normalization, to assess the proportion of genome-wide CpGs covered in each sample and identification of differential coverage between different contrasts. We used the package’s default parameters, as we have previously described (Pértille et al., 2017), except for the windows size parameter in which we mapped the CpN positions of the chicken reference genome to be used as region of interest (ROI), and also the chromosome selection parameter, of which we used only the chicken mtDNA. To map the CpNs we used the matchPattern function from “stringr” package from R. BSgenome.Ggallus.UCSC.galGal6 package from Bioconductor repository was uploaded as the reference of the chicken mitochondrial genome. After that, we create a function in R to merge, every single CpN tested for differential methylation with its respective mt annotation coordinates obtained from the Variant Effect Predictor (VEP) tool v.71 (McLaren et al., 2010). This function used part of the script available at https://github.com/stephenturner/solarplot/blob/master/solarplot.R. However, instead of the *solarplot,* we used plot functions in the R environment to show the differential methylation statistics distributed across the contrasts we have tested.

### DNA methylation analysis and functional annotation

For the 12 individuals, an average “enrichment score” of 2.2 (±0.1) was obtained to test the CpGs enrichment within the genomic regions covered by the given set of short reads compared to the full reference genome for which the “enrichment score” was 1.1 (table S11). This score was obtained dividing the relative frequency of CpGs within the given regions by the reference genome sequences. Non-enriched DNA sequences should present values close to 1, in contrast, a MeDIP-Seq experiment should return CpG rich sequences what will be indicated by increased CpG enrichment scores (Lienhard et al., 2014).

We then performed a differential methylation analysis focusing on CpNs. Our method interrogated differential coverage between the contrasts, individually, for each one of the CpN sites present on the chicken mtDNA genome that is 16,775 bps wide. For each site, we tested four contrasts: two comparing control vs stressed groups for male and female separately (MC vs MS and FC vs FS, respectively) and two to test for sex differences between the control and stressed groups (MC vs FC and MS vs FS). As we have described in the past (Pértille et al., 2017), epigenetic variations may be subtle or greater, depending on the environmental effect, so is worth to consider the use of multiple test corrections according to our findings. In this sense, we performed several tests and comparisons to provide a broad range of DMCpNs using a P-value threshold of 0.05 (Data S1). Significant DMCpNs were then annotated against the chicken reference genome *Gallus gallus* 6.0 (NCBI).

### Identifying methylation-dense regions

Upon visualising the significant dinucleotides obtained through our contrasts, we found that significantly differentially methylated cytosines clustered in distinct regions within the mtDNA (mtDMRs, supplementary table S2). In order to identify such regions, we constructed a model in which an mtDMR was defined by having a maximum gap of length x base pairs between adjacent differentially methylated cytosine motifs (CpNs). Through checking the number of merged windows generated with gap lengths, x ranging from 1-1962, we identified four important x-value ranges (127-179, 210-284,389-522 and 659-882). Within each range, increasing the gap length did not alter the number of merged regions. From 127 to 179, we identified the same 21 regions, giving a consecutive index (CI=53, representing the range of x for which the number of significant regions remains constant), while from 210 to 284, we identified 16 merged windows (CI = 76), from 389 to 522 we identified 11 windows (CI=124), and finally from 659 to 882, we identified 6 windows (CI= 223). High CI indicates low specificity because large changes in gap length do not change the number of dinucleotides which belong to that specific region. On the other hand, a low CI tends towards the border of significance among dinucleotides. We therefore used the smallest gap length (x = 127) that gave us the lowest CI, giving a conservative estimate of the number of mtDMRs.

### Motif discovery and enrichment analysis

In order to identify motifs within mtDMRs as well as elucidate their functionality, 46 of the of the methylation-enriched regions (5-387 bp in length) were submitted to XSTREME motif discovery and enrichment analysis, which both detects enriched motifs and then searches for homologues within vertebrate databases (version 5.5.0 (*60*), E values ≤ 0.05). Having identified enriched motifs within our mtDMRs and any vertebrate homologues within them, the sequence of homologous motifs were then drawn from human databases (for motifs, see tables S6, S7 and S8) and the occurrences of these sequences were determined within the mitochondrial genomes of *Gallus gallus, Homo sapiens, Alligator mississippiensis* and *Danio rerio* using FIMO (version 5.5.0 (*22*), p < 0.0001).

## Acknowledgments

The authors would like to thank the following funding agencies for their contributions to the research presented:

Formas 2019-02084 (PJ) and 2018-01074 (CGB)

European Research Council Advanced grant 322206 (“Genewell”) (PJ)

The Swedish Research council (VR) 2019-04053_VR (CGB, JL)

The São Paulo Research Foundation (FAPESP) projects #2016/20440-3 and #2018/13600-0 (FP)

## Author contributions

Experimental design: PJ

Experimental work and sample collection: PL

Conceptualization: JL, FP, CGB

Investigation: JL, FP, CGB Visualization: JL, FP Supervision: CGB, PJ

Writing—original draft: JL, FP

Writing—review & editing: PJ, CGP, FP, PL, JL

## Data and materials availability

All data needed to evaluate the conclusions in the paper are present in the paper and/or the Supplementary Materials

## References

1. M. G. P. van der Wijst, M. G. Rots, Mitochondrial epigenetics: an overlooked layer of regulation? Trends Genet. 31, 353–356 (2015).

2. M. Devall, J. Roubroeks, J. Mill, M. Weedon, K. Lunnon, Epigenetic regulation of mitochondrial function in neurodegenerative disease: New insights from advances in genomic technologies. Neurosci. Lett. 625, 47–55 (2016).

3. S. Dimauro, G. Davidzon, Mitochondrial DNA and disease. Ann Med 37, 222–232 (2005).

4. M. Picard, D. M. Turnbull, Linking the Metabolic State and Mitochondrial DNA in Chronic Disease, Health, and Aging. Diabetes 62, 672–678 (2013).

5. L. Lambertini, H. M. Byun, Mitochondrial epigenetics and environmental exposure. Curr Environ Health Rep 3, 214–224 (2016).

6. A. Stoccoro, F. Coppedè, Mitochondrial dna methylation and human diseases. Int. J. Mol. Sci. 22, 1–27 (2021).

7. V. Iacobazzi, A. Castegna, V. Infantino, G. Andria, Mitochondrial DNA methylation as a next-generation biomarker and diagnostic tool. Mol. Genet. Metab. 110, 25–34 (2013).

8. B. G. Janssen, H. M. Byun, W. Gyselaers, W. Lefebvre, A. A. Baccarelli, T. S. Nawrot, Placental mitochondrial methylation and exposure to airborne particulate matter in the early life environment: An ENVIRONAGE birth cohort study. Epigenetics 10, 536–544 (2015).

9. J. E. J. Wolters, S. G. J. Van Breda, F. Caiment, S. M. Claessen, T. M. C. M. De Kok, J. C. S. Kleinjans, Nuclear and Mitochondrial DNA Methylation Patterns Induced by Valproic Acid in Human Hepatocytes. Chem. Res. Toxicol. 30, 1847–1854 (2017).

10. D. A. Armstrong, B. B. Green, B. A. Blair, D. J. Guerin, J. F. Litzky, N. R. Chavan, K. J. Pearson, C. J. Marsit, Maternal smoking during pregnancy is associated with mitochondrial DNA methylation. Environ. Epigen. 2, dvw020–dvw020 (2016).

11. Q. Liu, H. Li, L. Guo, Q. Chen, X. Gao, P. h. Li, N. Tang, X. Guo, F. Deng, S. Wu, Effects of short-term personal exposure to air pollution on platelet mitochondrial DNA methylation levels and the potential mitigation by L-arginine supplementation. J. Hazard. Mater. 417, 125963–125963 (2021).

12. N. Fatima, S. Rana, Metabolic implications of circadian disruption. Pflueg. Arch. Eur. J. Physiol. 472, 513–526 (2020).

13. C. H. Ko, J. S. Takahashi, Molecular components of the mammalian circadian clock. Hum. Mol. Genet. 15 Spec No 2, R271–277 (2006).

14. V. M. Cassone, V. Kumar, “Circadian Rhythms” in Sturkie’s avian physiology (sixth edition), C. G. Scanes, Ed. (Academic Press, (2015)).

15. S. P. Karaganis, P. A. Bartell, V. R. Shende, A. F. Moore, V. M. Cassone, Modulation of metabolic and clock gene mRNA rhythms by pineal and retinal circadian oscillators. Gen. Comp. Endocrinol. 161, 179–192 (2009).

16. J. J. Gooley, K. Chamberlain, K. A. Smith, S. B. Khalsa, S. M. Rajaratnam, E. Van Reen, J. M. Zeitzer, C. A. Czeisler, S. W. Lockley, Exposure to room light before bedtime suppresses melatonin onset and shortens melatonin duration in humans. J. Clin. Endocrinol. Metab. 96, E463–472 (2011).

17. S. Moaraf, Y. Vistoropsky, T. Pozner, R. Heiblum, M. Okuliarová, M. Zeman, A. Barnea, Artificial light at night affects brain plasticity and melatonin in birds. Neurosci. Lett. 716, 134639 (2020).

18. S. A. Plano, L. P. Casiraghi, P. García Moro, N. Paladino, D. A. Golombek, J. J. Chiesa, Circadian and Metabolic Effects of Light: Implications in Weight Homeostasis and Health. Front. Neurol. 8, (2017).

19. D.-X. Tan, L. C. Manchester, X. Liu, S. A. Rosales-Corral, D. Acuna-Castroviejo, R. J. Reiter, Mitochondria and chloroplasts as the original sites of melatonin synthesis: a hypothesis related to melatonin’s primary function and evolution in eukaryotes. J. Pineal Res. 54, 127–138 (2013).

20. L. G. de Almeida Chuffa, F. R. F. Seiva, M. S. Cucielo, H. S. Silveira, R. J. Reiter, L. A. Lupi, Mitochondrial functions and melatonin: a tour of the reproductive cancers. Cell. Mol. Life Sci. 76, 837–863 (2019).

21. J. Cedernaes, M. Schönke, J. O. Westholm, J. Mi, A. Chibalin, S. Voisin, M. Osler, H. Vogel, K. Hörnaeus, S. L. Dickson, S. B. Lind, J. Bergquist, H. B. Schiöth, J. R. Zierath, C. Benedict, Acute sleep loss results in tissue-specific alterations in genome-wide DNA methylation state and metabolic fuel utilization in humans. Sci. Adv. 4, eaar8590 (2018).

22. C. E. Grant, T. L. Bailey, W. S. Noble, FIMO: scanning for occurrences of a given motif. Bioinformatics 27, 1017–1018 (2011).

23. D. M. Dominoni, J. C. Borniger, R. J. Nelson, Light at night, clocks and health: from humans to wild organisms. Biol. Lett. 12, 20160015 (2016).

24. E. L. Bittman, T. S. Kilduff, L. J. Kriegsfeld, R. Szymusiak, L. A. Toth, F. W. Turek, Animal care practices in experiments on biological rhythms and sleep: report of the Joint Task Force of the Society for Research on Biological Rhythms and the Sleep Research Society. J. Am. Assoc. Lab. Anim. Sci. 52, 437–443 (2013).

25. M. Mechta, L. R. Ingerslev, O. Fabre, M. Picard, R. Barrès, Evidence suggesting absence of mitochondrial DNA methylation. Front. Gen. 8, 1–9 (2017).

26. E. E. Hong, C. Y. Okitsu, A. D. Smith, C.-L. Hsieh, Regionally Specific and Genome-Wide Analyses Conclusively Demonstrate the Absence of CpG Methylation in Human Mitochondrial DNA. Mol. Cell. Biol. 33, 2683–2690 (2013).

27. S. Matsuda, T. Yasukawa, Y. Sakaguchi, K. Ichiyanagi, M. Unoki, K. Gotoh, K. Fukuda, H. Sasaki, T. Suzuki, D. Kang, Accurate estimation of 5-methylcytosine in mammalian mitochondrial DNA. Sci. Rep. 8, 5801 (2018).

28. R. Guitton, G. S. Nido, C. Tzoulis, No evidence of extensive non-CpG methylation in mtDNA. Nucleic. Acids. Res. 50, 9190–9194 (2022).

29. G. K. Marinov, Y. E. Wang, D. Chan, B. J. Wold, Evidence for Site-Specific Occupancy of the Mitochondrial Genome by Nuclear Transcription Factors. PLOS ONE 9, e84713 (2014).

30. M. Gardiner-Garden, M. Frommer, CpG islands in vertebrate genomes. J. Mol. Biol. 196, 261–282 (1987).

31. Y. Jia, R. Li, R. Cong, X. Yang, Q. Sun, N. Parvizi, R. Zhao, Maternal Low-Protein Diet Affects Epigenetic Regulation of Hepatic Mitochondrial DNA Transcription in a Sex-Specific Manner in Newborn Piglets Associated with GR Binding to Its Promoter. PLoS ONE 8, (2013).

32. A. Stoccoro, G. Siciliano, L. Migliore, F. Coppedè, Decreased Methylation of the Mitochondrial D-Loop Region in Late-Onset Alzheimer’s Disease. J. Alzheimer’s Dis. 59, 559–564 (2017).

33. H. Tong, L. Zhang, J. Gao, S. Wen, H. Zhou, S. Feng, Methylation of mitochondrial DNA displacement loop region regulates mitochondrial copy number in colorectal cancer. Mol. Med. Rep. 16, 5347–5353 (2017).

34. L. D. Zheng, L. E. Linarelli, L. Liu, S. S. Wall, M. H. Greenawald, R. W. Seidel, P. A. Estabrooks, F. A. Almeida, Z. Cheng, Insulin resistance is associated with epigenetic and genetic regulation of mitochondrial DNA in obese humans. Clin. Epigenet. 7, 1–9 (2015).

35. M. Mishra, R. A. Kowluru, Epigenetic modification of mitochondrial DNA in the development of diabetic retinopathy. Investig. Ophthalmol. Vis. Sci. 56, 5133–5142 (2015).

36. T. G. Demarest, M. M. McCarthy, Sex differences in mitochondrial (dys)function: Implications for neuroprotection. J. Bioenerg. Biomembr. 47, 173–188 (2015).

37. C. Bilu, N. Kronfeld-Schor, P. Zimmet, H. Einat, Sex differences in the response to circadian disruption in diurnal sand rats. Chronobiol. Int. 39, 169–185 (2022).

38. J. Qian, C. J. Morris, R. Caputo, W. Wang, M. Garaulet, F. Scheer, Sex differences in the circadian misalignment effects on energy regulation. Proc. Natl. Acad. Sci. U. S. A. 116, 23806–23812 (2019).

39. D. Bellizzi, P. D’Aquila, T. Scafone, M. Giordano, V. Riso, A. Riccio, G. Passarino, The control region of mitochondrial DNA shows an unusual CpG and non-CpG methylation pattern. DNA Res. 20, 537–547 (2013).

40. A. Nedoluzhko, R. Mjelle, M. Renström, K. H. Skjærven, F. Piferrer, J. M. O. Fernandes, The first mitochondrial 5-methylcytosine map in a non-model teleost (Oreochromis niloticus) reveals extensive strand-specific and non-CpG methylation. Genomics 113, 3050–3057 (2021).

41. V. Patil, C. Cuenin, F. Chung, J. R. R. Aguilera, N. Fernandez-Jimenez, I. Romero-Garmendia, J. R. Bilbao, V. Cahais, J. Rothwell, Z. Herceg, Human mitochondrial DNA is extensively methylated in a non-CpG context. Nucleic. Acids. Res. 47, 10072–10085 (2019).

42. J. E. Lee, M. Oney, K. Frizzell, N. Phadnis, J. Hollien, Drosophila melanogaster activating transcription factor 4 regulates glycolysis during endoplasmic reticulum stress. G3 (Bethesda) 5, 667–675 (2015).

43. A. Cagan, A. Baez-Ortega, N. Brzozowska, F. Abascal, T. H. H. Coorens, M. A. Sanders, A. R. J. Lawson, L. M. R. Harvey, S. Bhosle, D. Jones, R. E. Alcantara, T. M. Butler, Y. Hooks, K. Roberts, E. Anderson, S. Lunn, E. Flach, S. Spiro, I. Januszczak, E. Wrigglesworth, H. Jenkins, T. Dallas, N. Masters, M. W. Perkins, R. Deaville, M. Druce, R. Bogeska, M. D. Milsom, B. Neumann, F. Gorman, F. Constantino-Casas, L. Peachey, D. Bochynska, E. S. J. Smith, M. Gerstung, P. J. Campbell, E. P. Murchison, M. R. Stratton, I. Martincorena, Somatic mutation rates scale with lifespan across mammals. Nature 604, (2022).

44. J. H. Sun, S. M. Ai, S. Q. Liu, Methylation-driven model for analysis of dinucleotide evolution in genomes. Theor. Biol. Med. Model. 17, 3 (2020).

45. K. Sasaki, T. Uchiumi, T. Toshima, M. Yagi, Y. Do, H. Hirai, K. Igami, K. Gotoh, D. Kang, Mitochondrial translation inhibition triggers ATF4 activation, leading to integrated stress response but not to mitochondrial unfolded protein response. Biosci. Rep. 40, 1–12 (2020).

46. P. M. Quirós, M. A. Prado, N. Zamboni, D. D’Amico, R. W. Williams, D. Finley, S. P. Gygi, J. Auwerx, Multi–omics analysis identifies ATF4 as a key regulator of the mitochondrial stress response in mammals. J. Cell Biol. 216, 2027–2045 (2017).

47. K. R. Nitta, A. Jolma, Y. Yin, E. Morgunova, T. Kivioja, J. Akhtar, K. Hens, J. Toivonen, B. Deplancke, E. E. M. Furlong, J. Taipale, Conservation of transcription factor binding specificities across 600 million years of bilateria evolution. eLife 4, e04837 (2015).

48. W. E. Barry, C. S. Thummel, The Drosophila HNF4 nuclear receptor promotes glucose-stimulated insulin secretion and mitochondrial function in adults. eLife 5, (2016).

49. L. Palanker, J. M. Tennessen, G. Lam, C. S. Thummel, Drosophila HNF4 regulates lipid mobilization and beta-oxidation. Cell Metab. 9, 228–239 (2009).

50. L. Yang, Z. Chen, E. S. Stout, F. Delerue, L. M. Ittner, M. R. Wilkins, K. G. R. Quinlan, M. Crossley, Methylation of a CGATA element inhibits binding and regulation by GATA-1. Nat. Commun. 11, 2560 (2020).

51. J. Yang, J. R. Horton, D. Wang, R. Ren, J. Li, D. Sun, Y. Huang, X. Zhang, R. M. Blumenthal, X. Cheng, Structural basis for effects of CpA modifications on C/EBPβ binding of DNA. Nucleic. Acids. Res. 47, 1774–1785 (2019).

52. H. Hashimoto, D. Wang, J. R. Horton, X. Zhang, V. G. Corces, X. Cheng, Structural Basis for the Versatile and Methylation-Dependent Binding of CTCF to DNA. Mol. Cell 66, 711–720.e713 (2017).

53. R. Lister, M. Pelizzola, R. H. Dowen, R. D. Hawkins, G. Hon, J. Tonti-Filippini, J. R. Nery, L. Lee, Z. Ye, Q.-M. Ngo, L. Edsall, J. Antosiewicz-Bourget, R. Stewart, V. Ruotti, A. H. Millar, J. A. Thomson, B. Ren, J. R. Ecker, Human DNA methylomes at base resolution show widespread epigenomic differences. Nature 462, 315–322 (2009).

54. M. J. Ziller, F. Müller, J. Liao, Y. Zhang, H. Gu, C. Bock, P. Boyle, C. B. Epstein, B. E. Bernstein, T. Lengauer, A. Gnirke, A. Meissner, Genomic distribution and inter-sample variation of non-CpG methylation across human cell types. PLoS Genet 7, e1002389 (2011).

55. R. Landick, Transcriptional Pausing as a Mediator of Bacterial Gene Regulation. Annu. Rev. Microbiol. 75, 291–314 (2021).

56. G. Barshad, S. Marom, T. Cohen, D. Mishmar, Mitochondrial DNA Transcription and Its Regulation: An Evolutionary Perspective. Trends Genet. 34, 682–692 (2018).

57. Y. Yue, L. Ren, C. Zhang, K. Miao, K. Tan, Q. Yang, Mitochondrial genome undergoes de novo DNA methylation that protects mtDNA against oxidative damage during the peri-implantation window. Proc. Natl. Acad. Sci., 1–9 (2022).

58. H. Li, B. Handsaker, A. Wysoker, T. Fennell, J. Ruan, N. Homer, G. Marth, G. Abecasis, R. Durbin, The Sequence Alignment/Map format and SAMtools. Bioinformatics 25, 2078–2079 (2009).

59. M. Devall, R. G. Smith, A. Jeffries, E. Hannon, M. N. Davies, L. Schalkwyk, J. Mill, M. Weedon, K. Lunnon, Regional differences in mitochondrial DNA methylation in human post-mortem brain tissue. Clin. Epigenet. 9, 1–15 (2017).

60. C. E. Grant, T. L. Bailey, XSTREME: Comprehensive motif analysis of biological sequence datasets. bioRxiv, 2021.2009.2002.458722 (2021).

